# ACE-2 Blockade & TMPRSS2 Inhibition Mitigate SARS-CoV-2 Severity Following Cigarette Smoke Exposure in Airway Epithelial Cells In Vitro

**DOI:** 10.1101/2024.06.23.600238

**Authors:** Shah S Hussain, Emily Falk Libby, Jacelyn E Peabody Lever, Jennifer L Tipper, Scott E Phillips, Marina Mazur, Qian Li, Javier Campos-Gómez, Kevin S Harrod, Steven M Rowe

## Abstract

Cigarette smoking is associated with COVID-19 prevalence and severity, but the mechanistic basis for how smoking alters SARS-CoV-2 pathogenesis is unknown. A potential explanation is that smoking alters the expression of the SARS-CoV-2 cellular receptor and point of entry, angiotensin converting enzyme-2 (ACE-2), and its cofactors including transmembrane protease serine 2 (TMPRSS2). We investigated the impact of cigarette smoking on the expression of ACE-2, TMPRSS2, and other known cofactors of SARS-CoV-2 infection and the resultant effects on infection severity in vitro. Cigarette smoke extract (CSE) exposure increased ACE-2 and TMPRSS2 mRNA expression compared to air control in ferret airway cells, Calu-3 cells, and primary human bronchial epithelial (HBE) cells derived from normal and COPD donors. CSE-exposed ferret airway cells inoculated with SARS-CoV-2 had a significantly higher intracellular viral load versus vehicle-exposed cells. Likewise, CSE-exposure increased both SARS-CoV-2 intracellular viral load and viral replication in both normal and COPD HBE cells over vehicle control. Apoptosis was increased in CSE-exposed, SARS-CoV-2-infected HBE cells. Knockdown of ACE-2 via an antisense oligonucleotide (ASO) reduced SARS-CoV-2 viral load and infection in CSE-exposed ferret airway cells that was augmented by co-administration of camostat mesylate to block TMPRSS2 activity. Smoking increases SARS-CoV-2 infection via upregulation of ACE2 and TMPRSS2.

## INTRODUCTION

Coronavirus disease 2019 (COVID-19) was declared a global pandemic by the World Health Organization and is responsible for a devastating and ongoing mortality and morbidity burden worldwide. COVID-19 is caused by severe acute respiratory syndrome coronavirus 2 (SARS-CoV-2), a new member of the betacoronavirus genus that, like the betacoronavirus severe acute respiratory syndrome coronavirus (SARS-CoV) that emerged in humans in 2002, exploits the transmembrane protein angiotensin-converting enzyme-2 (ACE-2) as its key point of access into host cells (Hoffmann et al., 2020b; Lan et al., 2020). Cellular entry of SARS-CoV-2 requires the attachment of its spike (S) protein to the ACE-2 receptor via the S1 surface unit, in addition to the fusion of virus and cell membranes, accomplished by cleavage of the S protein at the S1/S2 and the S2’ site by the serine protease, transmembrane protease serine 2 (TMPRSS2) (Hoffmann et al., 2020a). ACE-2, which normally plays a role in protecting against pulmonary and cardiovascular injury via regulation of the renin-angiotensin system (Bourgonje et al., 2020; Gheblawi et al., 2020; Imai et al., 2005), and TMPRSS2 are broadly and heterogeneously expressed in human tissues, including within the respiratory tract (Baughn et al., 2020; Bourgonje et al., 2020; Ortiz Bezara et al., 2020; Sungnak et al., 2020). Noting the prominence of ACE-2 and TMPRSS2 in mediating the foothold of SARS-CoV-2 infection, it has been postulated that elevated ACE-2 and TMPRSS2 are a mechanistic link underlying greater COVID-19 infection rates and poorer outcomes among individuals known to be at high risk, although this relationship remains to be well elucidated in controlled models.

The clinical course of COVID-19 varies widely, in severe cases leading to critical illness from pneumonia and multiorgan failure that may result in death. Disease incidence and severity is associated with a spectrum of established risk factors, including underlying pulmonary conditions such as chronic obstructive pulmonary disease (COPD), cardiovascular disease, obesity, diabetes mellitus, immunosuppression, and older age (Berlin et al., 2020; Guan et al., 2020b; Sanyaolu et al., 2020). Cigarette smoking is among the additional factors that have been explored as a determinant of COVID-19 outcome, given the well accepted role of smoking in promoting lung injury and chronic lung disease (Centers for Disease et al., 2010). Meta-analyses conducted by the World Health Organization and others have revealed an association between smoking and heightened disease severity and death in people hospitalized with COVID-19 (https://www.who.int/news-room/commentaries/detail/smoking-and-covid-19), as well as an increased risk of severe COVID-19 disease progression in smokers, former smokers, and smokers with COPD (Patanavanich and Glantz, 2020). Similarly, COPD patients, for whom the majority of disease is due to cigarette smoking, have high incidences of COVID-19 severity and mortality as compared to the healthy population (Alqahtani et al., 2020; Emami et al., 2020; Zhang et al., 2020b). An investigation based on the National Health Interview Survey data implicated smoking as the most influential factor predisposing young adults to severe COVID-19 (Adams et al., 2020). These findings are complemented by a recent study demonstrating that cigarette smoke exposure augments the number of primary human nonsmoker cells infected by SARS-CoV-2 (Purkayastha et al., 2020), although it should also be noted that some reports have described a low prevalence of smokers among COVID-19 patients (Goyal et al., 2020; Guan et al., 2020a; Rossato et al., 2020) and even indicated that nicotine may have a protective effect against hyper-inflammatory response to SARS-CoV-2 (Farsalinos et al., 2020b), reflecting the complexity of the relationship and exposing the likely inconsistencies inherent in collection of smoking history.

Intriguingly, multiple investigations have found that ACE-2 (Cai et al., 2020b; Leung et al., 2020a; Matusiak and Schürch, 2020; Saheb Sharif-Askari et al., 2020; Smith et al., 2020a; Yin et al., 2020; Zhang et al., 2020a) and TMPRSS2 (Matusiak and Schürch, 2020; Saheb Sharif-Askari et al., 2020; Yin et al., 2020) are elevated in the airways of smokers and smokers with COPD, pointing to a putative pathway that may illuminate susceptibility to severe COVID-19 among smokers. In the current study, we sought to probe this relationship by evaluating complementary and controlled experimental models of SARS-CoV-2 infection. Here, we exploited a ferret model of cigarette smoke induced COPD, coupled with the evaluation of primary human bronchial epithelial (HBE) cells derived from smokers with and without COPD. We evaluated ACE-2 and TMPRSS2 expression, in addition to other known cofactors of SARS-CoV-2 infection, in response to smoking, and consequent effects on infection severity. The results demonstrate evidence that smoking increases the risk and severity of COVID-19 infection through the combined induction of ACE-2 receptor expression and TMPRSS2, establishing the basis of increased severity of COVID-19 among smokers. Our studies further show that simultaneous blockade of both ACE-2 and TMPRSS2 pathways were required to ameliorate infection, suggesting therapeutic implications.

## RESULTS

### ACE-2 and TMPRSS2 expression are elevated in cigarette smoke-exposed ferret lung

Chronic cigarette smoke exposure is associated with the induced expression of mucin-expressing secretory cells, cell types that highly express ACE-2 and TMPRSS2 (Hussain et al., 2018; Johnson, 2011; Lukassen et al., 2020). We used a COPD ferret model of chronic cigarette smoke exposure to directly assess the effect of cigarette smoke on the expression of ACE-2, TMPRSS2, and other proteins associated with SARS-CoV-2 entry and replication (Zhang et al., 2020a). Ferrets were exposed to cigarette smoke or air control for six months, after which lung tissue was collected for analysis. Preliminary studies in smoke-exposed ferrets using RNA-Seq demonstrated that ACE-2 and TMPRSS2 were among the SARS-CoV-2 relevant pathways substantially elevated, among others (Fig. 1A). This observation was recapitulated by qRT-PCR analyses demonstrating significantly elevated ACE-2 (Fig. 1B: 1.4-fold, P=0.007) and TMPRSS2 (Fig. 1C: 1.8-fold, P=0.015) mRNA levels in smoked-exposed vs. air control tissue, as also visualized by immunohistochemical staining (Fig. 1D,E). To understand whether increased ACE-2 and TMPRSS2 expression were also present in large ferret airways, and thus more likely to be involved in SARS-CoV-2 acquisition and descending infection, we next exposed terminally differentiated normal FTECs to CSE (1%, apical) for 48 hr and repeated qRT-PCR. Indeed, CSE exposure resulted in an increase in ACE-2 (Fig. 1F: 1.4-fold, P=0.002) and TMPRSS2 (Fig. 1G: 2.0-fold, P=0.0006) mRNA expression over the vehicle. Overall, these data indicate that both chronic in vivo and acute in vitro cigarette smoke exposure increases the expression of ACE-2 and TMPRSS2 in ferret lung cells. Findings are consistent with a similar report of the human lung transcriptome (Jacobs et al., 2020; Leung et al., 2020b), but in an experimental system that controlled for covariates to human smoking not readily obtained in human studies.

**Figure 1.**
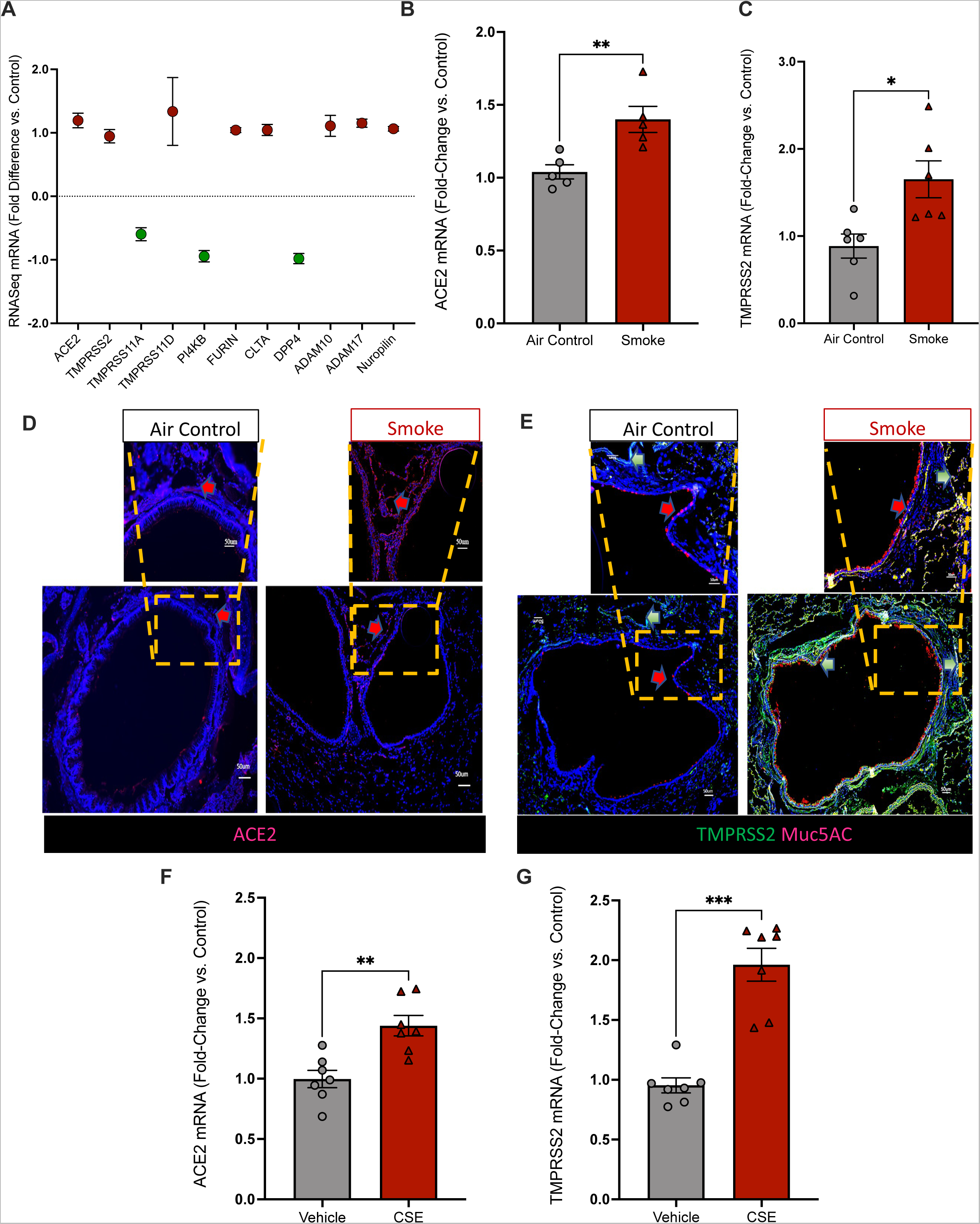
ACE-2 and TMPRSS2 expression are elevated in cigarette smoke-exposed ferret lung. (A) Dot plot showing from RNA-Seq analysis of SARS-CoV-2 receptor and associated gene expression in lung tissue of ferrets exposed to cigarette smoke for six months. (B) ACE-2 and (C) TMPRSS2 mRNA expression in lung tissue from smoke-exposed vs. air control ferrets as assessed by qRT-PCR.(D) Representative immunofluorescence images of smoke-exposed and air control ferret lung sections stained with ACE-22 antibody (red) and DAPI (blue). Scale bar,50uM. Areas of magnification (magnification 40X) are outlined by the red dashed line. .(E) Representative immunofluorescence images of smoke-exposed and air control ferret lung sections stained with TMPRSS-2 antibody (Green), Muc5AC (red), and DAPI (blue). Scale bar,50uM. Areas of magnification (magnification 40X) are outlined by the red dashed line qRT-PCR analysis of (F) ACE2 and (G) TMPRSS2 mRNA expression in terminally differentiated ferret tracheal epithelial cells (FTECs) exposed to cigarette smoke extract (CSE) or vehicle control. *P<0.05, **P<0.005,***P<0.0005.

### ACE-2 and TMPRSS2 expression are elevated in CSE-exposed human airway cells

To assess the effect of cigarette smoke on ACE-2 and TMPRSS2 expression in human cells, we proceeded with studies using the human lung Calu-3 cell line (a cell type that demonstrates sensitivity to SARS-CoV-2 infection (Li et al., 2020) and bronchial epithelial cells from both healthy donors (healthy HBE) and patients with COPD (COPD HBE). In Calu-3 cells, CSE exposure increased ACE-2 (Fig. 2A: 1.5-fold, P=0.002) and TMPRSS2 (Fig. 2B: 1.7-fold, P=0.002) mRNA expression compared to vehicle control. Correspondingly, in normal HBE cells, CSE exposure elicited 2.8-fold (P=0.0079; Fig. 2C) and 2.7-fold (P=0.02; Fig 2D) increases in ACE-2 and TMPRSS2 expression, respectively, compared to vehicle. Many effects of COPD evident in the airways of patients including mRNA expression profiles (Mathis et al., 2013; Pezzulo et al., 2011)—persist in the airway epithelial cells derived from these individuals. In HBE cells derived from COPD donors, mRNA expression of ACE-2 (Fig. 2E) and TMPRSS2 (Fig. 2F) was increased (ACE-2, 2.3-fold, P=0.04; TMPRSS2, 1.9-fold, P=0.0019 compared to HBE from healthy non-smoking controls. Western blot analysis revealed that TMPRSS2 protein expression was also significantly upregulated in COPD tissue (Fig. 2G,H). Overall, these results indicate that elevated ACE-2 and TMPRSS2 expression is evident with acute cigarette exposure in normal human airway cells and is also persistent in patients with COPD who once but no longer smoked, suggesting a propensity for more severe COVID-19 in these individuals.

**Figure 2.**
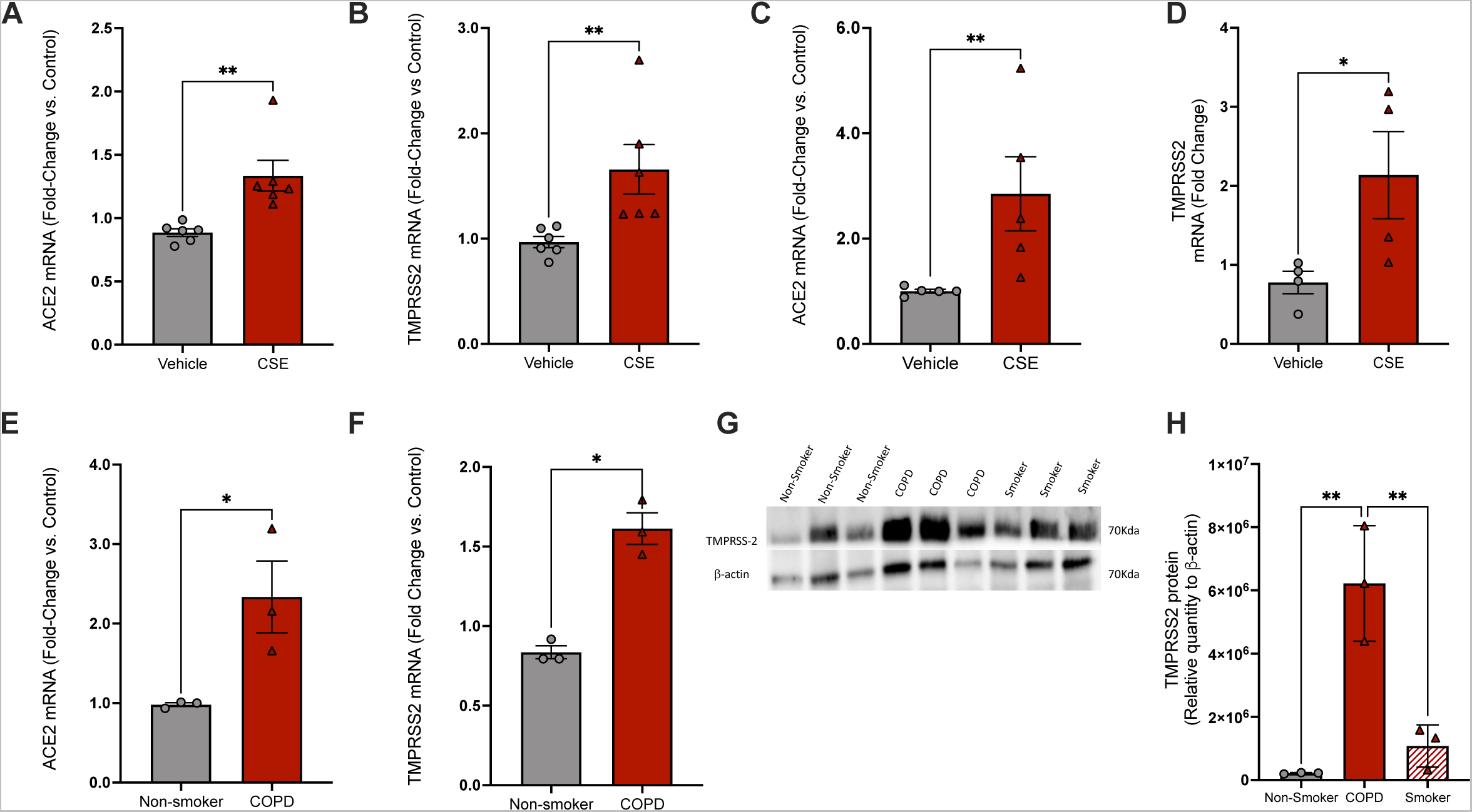
ACE-2 and TMPRSS2 expression are elevated in CSE-exposed human airway cells. qRT-PCR measurement of (A) ACE-2 and (B) TMPRSS2 mRNA expression in Calu-3 cells exposed to cigarette smoke extract (CSE) or vehicle control for 48 hr.qRT-PCR measurement of (C) ACE-2 and (D) TMPRSS2 mRNA expression in primary human bronchial epithelial (HBE) cells derived from healthy control donors and treated with CSE or vehicle control for 48 hr. N=3 monolayers/condition across 3-4 different donors. qRT-PCR measurement of (E) ACE-2 and (F) TMPRSS2 mRNA expression in bronchial epithelial cells derived from patients with COPD (COPD HBE) or healthy non-smoker donors. N=3 monolayers/condition across 3-4 different donors. (G) Representative western blot and (H) quantification of TMPRSS2 expression in COPD or healthy non-smoker donors. N = 3-4 monolayers/conditions derived from 3-4 different donors. *P<0.05, **P<0.005.

### SARS-CoV-2 infection is increased in CSE-exposed FTECs and Calu-3 cells

Increased expression of ACE-2 and TMPRSS2 observed with smoking indicated the potential for increased severity of SARS-CoV-2 infection. We therefore next designed studies of CSE exposure followed by inoculation with SARS-CoV-2 in the FTEC and Calu-3 cell models (Fig. 3A). In FTECs, SARS-CoV-2 genomic copy numbers were significantly higher after 72 hr of infection in CSE-treated (3.1 x 10^5^ ± 1.7 x 10^4^, P=0.006) vs. vehicle-treated (1.7 x 10^5^ ± 3.4 x 10^4^) cells (Fig. 3B). Consistent with this, significantly higher viral replication was evident from the apical media of CSE-treated FTECs, as determined by visualizing (Fig. 3D) and quantifying (Fig. 3C) foci forming units of Vero-E6 cells (CSE, 7.7 x 10^4^ ± 6.6 x 10^3^; vehicle, 5.6 x 10^4^ ± 6.0 x10^3^; P=0.048). Calu-3 cells exhibited a similar susceptibility to increased SARS-CoV-2 infection upon CSE exposure (Fig. 3E-G). These findings demonstrate that CSE increases the infectivity of ferret and human cells, further implicating elevated ACE-2 and TMPRSS2 expression as a putative pathway underlying the relationship.

**Figure 3.**
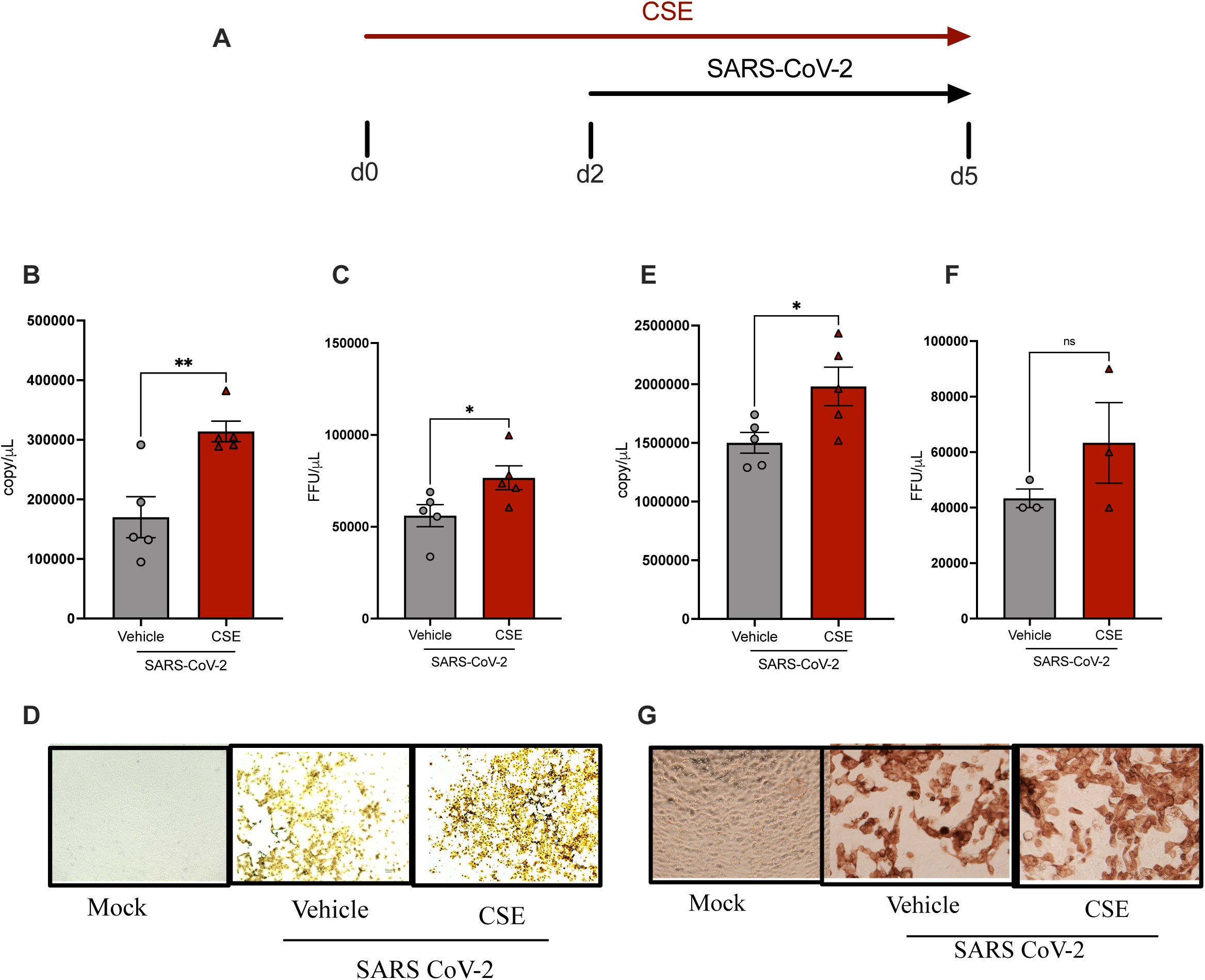
SARS-CoV-2 infection is increased in CSE-exposed FTECs and Calu-3 cells. (A) Schematic outline for experiments evaluating the relationship between cigarette smoking and SARS-CoV-2 infection in ferret tracheal epithelial cells (FTECs) and Calu-3 cells. Cells were treated with cigarette smoke extract (CSE) or vehicle control for 48 hr prior to inoculation with SARS-CoV-2 (MOI-3). After 72 hr days of SARS-CoV-2 infection with concomitant CSE exposure, cells were harvested for analysis. (B) Quantification of viral load by qRT-PCR of SARS-CoV-2 mRNA at three days post-infection in FTECs exposed to CSE or vehicle control, with (C) visualization and (D) quantification of SARS-CoV-2-infected cells by foci forming assay in Vero-E6 cells. N=3-5 well per conditions. (E) Quantification of viral load by qRT-PCR at three days post SARS-CoV-2 infection in Calu-3 cells exposed to CSE or vehicle control, with (F) visualization and (G) quantification of SARS-CoV-2-infected cells by foci forming assay in Vero-E6 cells. N=3-5 well per conditions. *P<0.05, **P<0.01.

### SARS-CoV-2 genomic replication is increased in CSE-exposed normal HBE cells and in HBE cells derived from COPD patients

Next, we examined SARS-CoV-2 replication in CSE-treated HBE cells derived from normal healthy controls as compared to COPD HBE cells. For consistency, the same cell donors were used in these studies as in the studies evaluating the effect of CSE on ACE-2 and TMPRSS2 levels (see Fig. 2C-F), and CSE exposure and SARS-CoV-2 infection periods remained constant from those depicted in Figure 3A. To evaluate epithelial integrity after inoculation with SARS-CoV-2 and prior to analysis, we obtained transepithelial electrical resistance (TEER) measurements (Fig. 4A). With SARS-CoV-2 infection, we observed a significant reduction in TEER in both CSE-treated normal HBE (17.27%) and COPD HBE (29.10 cells compared to vehicle-treated HBE cells, an effect not observed with mock infection. Mean SARS-CoV-2 copy number was also significantly higher in CSE-exposed (Fig. 4B: 1.3 x 10^5^ ± 1.1 x 10^5^, P=0.0038) vs. vehicle-exposed HBE (1.4 x 10^6^ ± 7.9 x 10^5^), as also seen in COPD HBE (Fig. 4C: COPD ; 3.5x 10^7^ ± 2.8 x 10^7^; non-smoker, 1.8X10^5^± 9.7 x 10^4,^ P=0.021, in line with our findings in FTECs and Calu-3 cells. SARS-CoV-2 infection tracked with ACE-2 and TMRSS2 expression, as visualized by RNAScope in SARS-CoV-2-infected (Fig. 4E) but not mock-infected (Fig. 4D) cells, although cells from COPD donors had more diffuse, rather than discrete SARS-CoV-2 replication along the monolayer.

**Figure 4.**
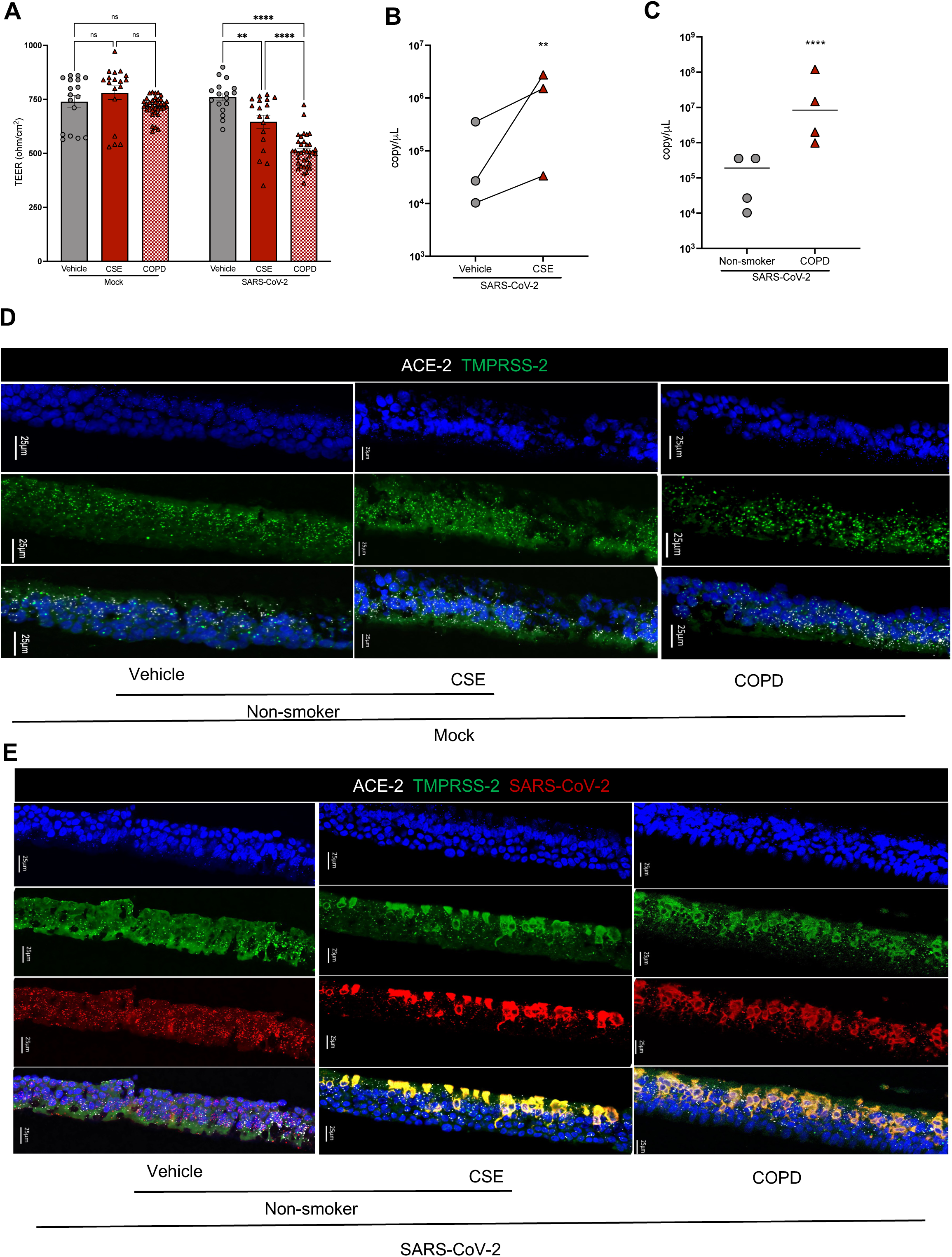
SARS-CoV-2 genomic replication is increased in CSE-exposed normal HBE cells and in HBE cells derived from COPD patients. (A) Transepithelial electrical (TEER) measurements in vehicle-treated, cigarette smoke extract (CSE)-treated, and COPD human bronchial epithelial (HBE) monolayer cultures infected with SARS-CoV-2 or mock infection. The data were obtained by averaging three independent Transwell reads, each of which represented the mean of three separate readings. (N=3-5) HBE. Each line corresponds to a distinct donor (3-5 donors in total), and each donor is represented by 3-5 replicates. (B) Quantification of viral load by qRT-PCR of SARS-CoV-2 mRNA at 72 hr post-infection with SARS-CoV-2 in CSE- or vehicle-exposed HBE cells. N=3 different donors, with each line representing a separate donor. (C) Quantification of viral load by qRT-PCR of SARS-CoV-2 mRNA at 72 hr post infection with SARS-CoV-2 in healthy non-smoker or COPD HBE cells. N= monolayers/condition derived from 3-5 different donors. (D-E) Representative images using RNAscope in situ hybridization for comparable detection of genomic RNA with only the SARS-CoV-2-specific S probe. Images show CSE- or vehicle-exposed HBE cells, and healthy non-smoker or COPD HBE cells, at 72 hr post inoculation with (D) mock infection or (E) SARS-CoV-2. SARS-CoV-2 (red), TMPRSS2 (green), ACE-2 (white), Nuclei (blue). **P<0.01.

We further evaluated smoking-related factors that may contribute to SARS-CoV-2 replication. First, to confirm the presence of mucin-expressing secretory cells, which are induced by cigarette smoke (Hussain et al., 2018; Johnson, 2011) and have been reported by some (Lukassen et al., 2020; Sungnak et al., 2020; Ziegler et al., 2020), but not all (Lee et al., 2020), to highly express ACE-2 and TMPRSS2, we proceeded with histopathological assessment (Supplemental Fig. 1A). Indeed, increases in Alcian Blue with periodic acid and Schiff’s solution (AB-PAS) staining consistent with mucus expression were observed in CSE-exposed normal HBE and COPD HBE compared to controls. With SARS-CoV-2 infection, the proportion of mucus-expressing cells did not meaningfully change in this time frame. In corroboration, key transcription factors associated with goblet cell hyperplasia from smoking— SAM-pointed domain-containing Ets-like factor (SPDEF) and forkhead box A3 (FOXA3), which cause goblet cell differentiation (Chand et al., 2018; Park et al., 2007; Rajavelu et al., 2015), and Forkhead box A2 (FOXA2), an inhibitor of goblet cell differentiation (Tang et al., 2013)— were elevated or decreased, respectively, regardless of whether SARS-CoV-2 was present or not (Supplemental Fig. 1C-G). Elevated expression of mucins MUC5AC and MUC5B and trefoil factor 3 (TFF3), factors associated with increased mucus from smoking, was also observed generally irrespective of SARS-CoV-2 infection, with the exception of TFF3 for which activation of expression from SARS-CoV-2 was also observed (Supplemental Fig. 1H-N).

We also explored whether antecedent epithelial injury induced by smoking (Heijink et al., 2012; Kosmider et al., 2011; van der Toorn et al., 2009) may play a role in facilitating SARS-CoV-2 induced cell injury. As seen in the representative images of Terminal deoxynucleotidyl transferase (TdT) dUTP Nick-End Labeling (TUNEL) staining in normal HBE cells upon CSE exposure and SARS-CoV-2 infection (Supplemental Fig. 2A), CSE markedly increased the percentage of cells expressing the apoptotic marker in response to SARS-CoV-2 infection (CSE, 60.4 ± 0.6 vs. vehicle, 25.0 ± 1.6; P<0.0001), while CSE also increased TUNEL in the absence of infection (Supplemental Fig. 2B: CSE, 20.3 ± 2.6 vs. vehicle, 4.3 ± 1.3; P<0.001), as expected, but to a lesser extent.

Together, these results suggest that cigarette smoke exposure and individuals with COPD have an increased susceptibility to SARS-CoV-2 infection compared to healthy controls in vitro, and this corresponds to increased ACE-2 and TMPRSS2 expression. The data also indicate that cigarette smoke predisposes the epithelium to injury from SARS-CoV-2, plausibly providing an additional pathway beyond SARS-CoV-2 cellular entry that mediates SARS-CoV-2 severity.

### Simultaneous ACE-2 blockade and TMPRSS2 inhibition reduce SARS-CoV-2 infection after CSE exposure in FTECs

To further probe the contribution of ACE-2 and TMPRSS2 to cigarette smoking-related COVID-19 disease severity, we investigated whether ACE-2 and TMPRSS2 inhibition could protect against cigarette smoke-enhanced SARS-CoV-2 infection. For ACE-2 blockade, we screened five chemically modified ACE-2-targeting ASOs in FTECs, selecting the compound that most efficaciously inhibited ACE-2 mRNA expression for our studies (Supplemental Fig. 3A); this ASO exhibited a dose-dependent effect within 7 days of incubation, achieving significant knockdown at all concentrations tested (Supplemental Fig. 3B). In FTECs treated according to the scheme depicted in Figure 5A, we further found that ASO blockade (20 µM) was able to reduce ACE-2 expression from CSE exposure, both in the absence (ACE-2 ASO, 0.2 ± 0.0 vs. control ASO, 1.3 ± 0.1; P<0.0001) and presence (ACE-2 ASO, 0.3 ± 0.0 vs. control ASO, 1.3 ± 0.2; P<0.0001) of SARS-CoV-2 (Fig 5B), an effect also seen in vehicle-exposed cells (no virus: ACE-2 ASO, 0.1 ± 0.0 vs. control ASO, 1.0 ± 0.0; P<0.0001 and SARS-CoV-2: ACE-2 ASO, 0.4 ± 0.1 vs. control ASO, 0.95 ± 0.01; P=0.015). ACE-2 ASO also significantly reduced TMPRSS2 expression from CSE exposure in the absence of virus (ACE-2 ASO, 1.7 ± 0.1 vs. control ASO, 2.2 ± 0.0; P=0.021), consistent with a report indicating a positive correlation between ACE-2 and TMPRSS levels (Gkogkou et al., 2020), although did not have an appreciable impact with vehicle exposure or in the presence of SARS-CoV-2, possibly due to lower baseline levels beyond which blockade by ACE-2 ASO had modest additional effect (Fig. 5C).

**Figure 5.**
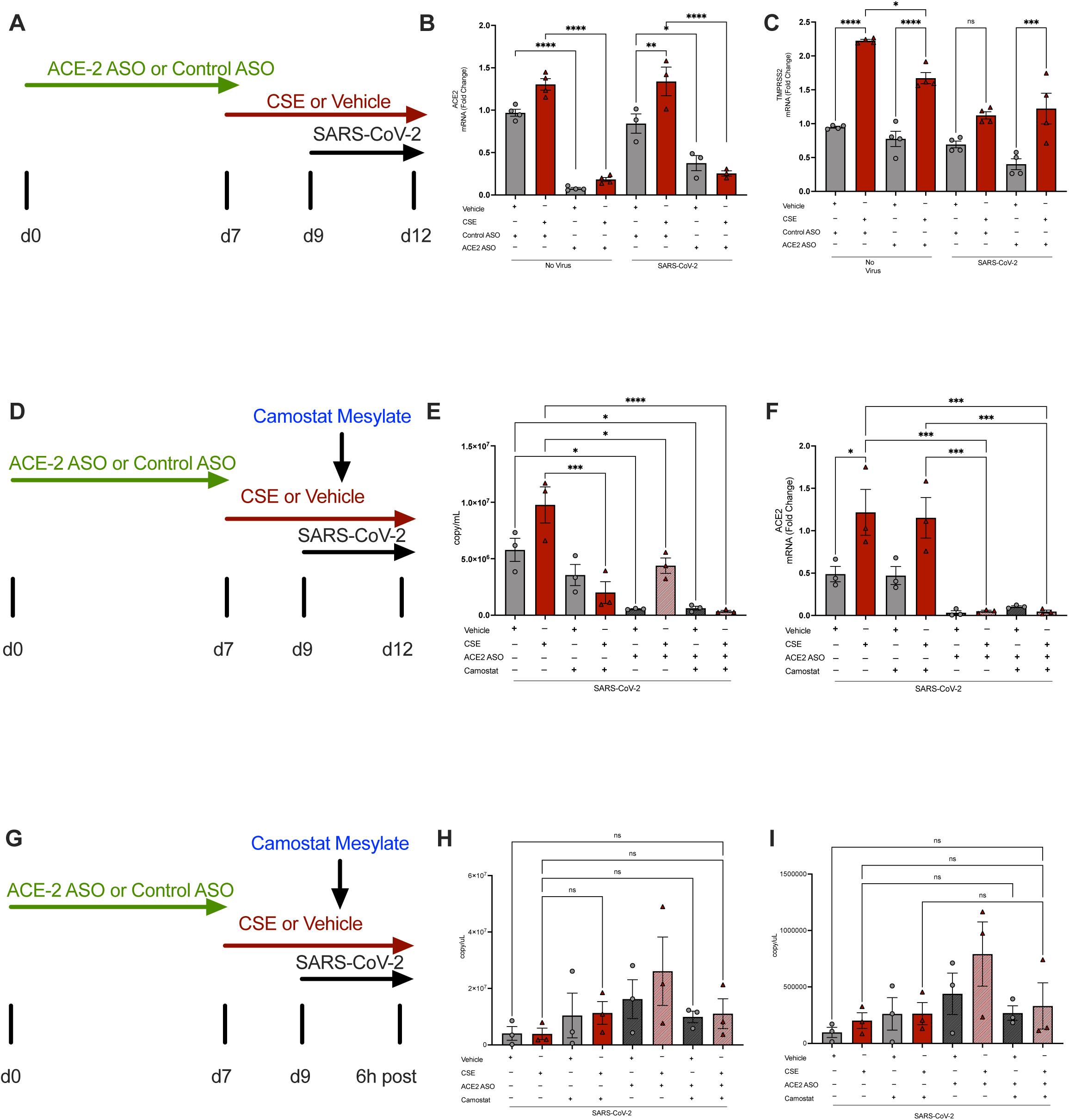
Simultaneous ACE2 blockade and TMPRSS2 inhibition reduces SARS-CoV-2 infection after CSE exposure in FTECs. (A) Scheme depicting the approach to ACE2 antisense oligonucleotide (ASO) treatment, cigarette smoke extract (CSE) exposure, and SARS-CoV-2 infection in ferret tracheal epithelial cells (FTECs). Seven days of treatment with ACE2 ASO (20 µM) or control ASO was followed by a 48-hr incubation period in CSE or vehicle control prior to inoculation with SARS-CoV-2 or no virus for 72 hr of infection. Assessment of mRNA expression of (B) ACE-2 and (C) TMPRSS2 following the scheme depicted in (A). (D) Scheme depicting the approach to ACE2 ASO and camostat mesylate treatment, CSE exposure, and SARS-CoV-2 infection in FTECs. Camostat mesylate (100 mM) was added for 2 hr on day 9, prior to inoculation with SARS-CoV-2. Assessment of (E) viral load and mRNA expression of (F) ACE-2 and (G) SARS-Cov2 infection for a shorter duration following the scheme depicted in (D). Viral load following shorter duration of infection (H) and (I). qRT-PCR was used for mRNA quantification in all studies. *P<0.05, **P<0.01, ***P<0.001, ****P<0.0001.

Based on these findings, we proceeded with examining the effect of ACE-2 ASO treatment alone and simultaneously with the pharmacological protease inhibitor camostat mesylate on SARS-CoV-2 infection in the FTEC model. Camostat mesylate is a broad spectrum protease inhibitor that has been administered to the nasal passage of human subjects for its properties as a channel activating protease inhibitor of the epithelial sodium channel, ENaC, and is presently under investigation in COVID-19 after showing inhibition of SARS-CoV-2 in vitro (Gunst et al., 2021; Hoffmann et al., 2020a). When administered to CSE-exposed cells as depicted by the study scheme in Figure 5D, both ACE-2 ASO (4.4 x 10^6^ ± 6.8 x 10^5^ vs. no treatment, 9.8 x 10^6^ ± 1.6 x 10^6^; P=0.012) and camostat mesylate (100 mM) (2.0 x 10^6^ ± 9.7 x 10^5^; P=0.0002) as single agents induced a significant reduction in SARS-CoV-2 viral copy number compared to untreated CSE-exposed cells that was augmented upon co-administration of the two approaches (ACE-2 ASO + camostat mesylate, 3.4 x 10^5^ ± 8.6 x 10^4^; P<0.0001) (Fig. 6E). Significant reductions in SARS-CoV-2 copy number were also observed in vehicle-exposed cells treated with ACE-2 ASO (5.5 x 10^5^ ± 3.1 x 10^4^ vs. no treatment, 5.8 x 10^6^ ± 1.0 x 10^6^; P=0.015) and, to a similar extent, combination ACE-2 ASO and camostat mesylate therapy (ACE-2 ASO + camostat mesylate, 6.2 x 10^5^ ± 1.7 x 10^5^; P=0.017). With CSE exposure, the impact of ACE-2 ASO and camostat mesylate coadministration on SARS-CoV-2 infection was accompanied by a pronounced downregulation of ACE-2 mRNA (ACE-2 ASO + camostat mesylate, 0.04 ± 0.0 vs. no treatment, 1.2 ± 0.3; P=0.001), but not more so than with ACE-2 ASO alone (0.05 ± 0.0; P=0.001), suggesting that additional co-receptors such as TMPRSS2 may compensate when ACE-2 expression is reduced (Fig. 5F). Interestingly, CSE-induced upregulation of MUC5AC was significantly attenuated by ACE-2 ASO (1.4 ± 0.2, vs. no treatment, 4.0 ± 0.0; P<0.0001), camostat mesylate (1.8 ± 0.0; P<0.0001), and to the greatest magnitude, coadministration of the two approaches (ACE-2 ASO + camostat mesylate, 0.3 ± 0.0; P<0.0001) (Supplemental Fig . 4B), implicating an interaction between SARS-CoV-2 entry factors, or SARS-CoV-2 infection itself, and smoking-related mucus expression. As opposed to the prominent effects seen at 72 hr post-infection, treatment with neither ACE-2 ASO nor camostat affected SARS-CoV-2 copy number when measured after 6 hr of infection (Fig. 5G), either by detection of genomic (Fig. 5H) or sub-genomic (Fig. 5I) primers. This suggests the principal effects of smoking were on replication rather than early events related to viral entry alone and also explains why inhibition of both viral entry and replication were required to exhibit benefit.

## DISCUSSION

It has been widely reported that the primary SARS-CoV-2 cell entry receptor, ACE-2, and cofactor, TMPRSS2, are elevated in the context of the smoking (Cai et al., 2020b; Leung et al., 2020a; Matusiak and Schürch, 2020; Saheb Sharif-Askari et al., 2020; Smith et al., 2020a; Yin et al., 2020; Zhang et al., 2020a), although the relationship between smoking and severe disease from SARS-CoV-2 infection has been a matter of debate. Here we have shown in ferret and human cell models that cigarette exposure induces upregulation of ACE-2 and TMPRSS2 that correlates with experimental augmentation of SARS-CoV-2 infection, a finding that also extended to airway cells from patients with COPD, which retain the injury signals conferred by smoking. These data provide direct evidence of a mechanistic pathway whereby smoking promotes ACE-2 receptor and TMPRSS2 expression, in turn enhancing SARS-CoV-2 cellular entry, increasing viral load, and conferring a greater risk of poor clinical outcome.

To date, studies of ACE-2 expression and Its correlation with SARS-CoV-2 infection in smokers and COPD is limited and contradictory (Papakonstantinou et al., 2015; Purkayastha et al., 2020). Some meta-analyses also suggest that smokers are not at risk of higher SARS-CoV-2 infection(Farsalinos et al., 2020a). Viral tropism has been reported that the SARS-COV-2 can infect and propagate differently in different regions of the body exposed to the virus. (Nakayama et al., 2021). While increased ACE-2 expression is a factor in viral entry, the molecules crucial for viral replication, particularly among smokers, remain uncertain. So, understanding of host-virus interaction will give novel insight to understand the infectivity and replication of different strains of the virus. We thus investigated the expression of ACE-2 and several associated proteases linked with SARS-CoV-2 processing in cells derived from smokers and non-smokers. Our data in vitro studies from ferret tracheal cells and human primary epithelial cells demonstrates that short exposure to CSE induces increased expression of the SARS-CoV-2 receptor ACE-2 and the protease TMPRSS2 necessary for SARS CoV2 cellular entry. These data are consistent with earlier reports where the increased expression of ACE-2 was observed by transcriptome analysis and single-cell sequencing (Cai et al., 2020a; Caruso et al., 2021; Smith et al., 2020b; Zhang et al., 2020a).

From our data dual blockade could be an explanation for the failed clinical trial of chemostat mesylate (NCT04353284). To provide further insight into other factors associated with SARS-CoV-2 entry and replication, we assessed the expression of TMPRSS2 and TMPRSS4, by qPCR. Interestingly, TMPRSS4 and Muc5AC, which have the ability to share fusion peptides in spike protein by proteolysis (Shulla et al., 2011), Upregulated mucin mRNA in case of smokers might have higher viral shedding in the lungs of smokers and were upregulated in the case of CSE-treated FTEC (Supplemental Figure 4). This aligns with the findings of Zhang et al. (2020), illustrating that various cell types possess receptors that could be linked to the infection. The expression of ACE-2 and other associated receptors differed among different cell types, rendering them susceptible to infection.(Zhang et al., 2020a)

Given the primacy of ACE-2 and TMPRSS2 in driving SARS-CoV-2 infection, efforts to develop therapies for patients with COVID-19 have naturally included a focus on modulation of ACE-2 (Jia et al., 2021; Monteil et al., 2020; Wang et al., 2021) and TMPRSS2 function (Chen et al., 2021; Hoffmann et al., 2020a; Leach et al., 2021; Maggio and Corsini, 2020), with a range of approaches reaching various stages of clinical development (Aldea et al., 2021; Gunst et al., 2021) (NCT04335136 ). We observed that an ASO targeting ACE-2 successfully reduced ACE-2 expression in a dose-dependent fashion, conferring a partial benefit on SARS-CoV-2 infection in ferret epithelial cells. Moreover, camostat mesylate demonstrated an ability to abrogate the deleterious effect of acute smoke exposure on viral load, and coadministration of camostat mesylate and ACE-2 ASO was most efficacious toward this end. That a TMPRSS2-targeting agent effectively reduced viral load in CSE-exposed cells alone and augmented the effect of ACE2 blockade suggests that co-receptors such as TMPRSS2 represent important pathways on their own. These co-receptors may also contribute a compensatory response when ACE2 expression is reduced. These findings suggest that ACE-2-directed strategies and pharmacological interventions targeting TMPRSS2 may be promising in COVID-19 patients when risk factors such as smoking are present. A dual therapy approach that modulates both ACE-2 and TMPRSS2 may thus be most beneficial. This echoes recently published studies reporting discordant findings on whether ACE-2 expression levels are correlated to SARS-CoV-2 infection severity, such as the infectious potential of b cells that can become infected despite low ACE-2 expression (Millette et al., 2021). Such findings are thought to influence b cell function in patients with diabetes mellitus (Wu et al., 2021)

Overall, these insights suggest that ACE2-centric strategies and pharmacological actions targeting TMPRSS2 might be valuable, especially for COVID-19 patients with risk factors like smoking. Consequently, a combined therapeutic approach addressing both ACE2 and TMPRSS2 might offer the highest benefit.

This could be explained by the distinction between viral entry, replication, and injury response. Smoke augments each of these prcesses. Our data suggests that entry and replication must both be addressed to meaningfully alter the increased risk of smoking to these pathways.

Additionally, it is conceivable that the injurious cellular effects of cigarette smoke accumulate, rendering cells more susceptible to SARS-CoV-2 infection and augmenting cellular responses including redox stress and induction of apoptosis. Like others (Purkayastha et al., 2020), we observed a heightened apoptotic response in the presence of smoke and SARS-CoV-2. It may be that this phenomenon is also associated with the propensity for diffuse cell necrosis and induction of cytokine storms seen in severe SARS-CoV-2 infections.

### Limitation of the Study

In summary, our data demonstrate that cigarette smoke exacerbates SARS-CoV-2 infection and resultant injury via more than one cellular entry mechanism. Given these multiplicitous pathways, combination therapies targeting various aspects of SARS-CoV-2 cellular entry may be needed in smokers to circumvent infection and severe COVID-19 in this population. In our study, we lack the study where we would like to activate the inhibition study of associated factors. However, other studies have shown the importance of these factors too. Some of the studies of ACE-2 blockade were not feasible in the case of HBE cells due to lack of ASOs that could efficiently knockdown ACE-2 as in the case of FBE,

## MATERIALS AND METHODS

### Ferret smoking

Wild-type ferrets (*Mustela putorius furo*, females, 0.6–0.8 kg in body weight; males, 1.2–2.0 kg in body weight) matched by age and sex were obtained from Marshall BioResources, subject to a brief acclimatization and training period, and randomized to receive air control or whole cigarette smoke exposure. Ferrets were then restrained in customized male and female nose-only exposure tubes that entailed a 24-port plenum connected to the mainstream smoke output. Using 1R6F research cigarettes, cigarette smoke was generated by an automated 700700 series cigarette smoke generator (TSE Systems; Chesterfield, MO) with a 10-cigarette carousel and built-in software calibration, and animals were exposed to 60 min of smoke, twice daily over a six-month period, as described. (Kaza et al., 2021; Raju et al., 2016) The accuracy and safety of exposure were monitored using an in-line gas analyzer for oxygen, carbon monoxide, and particulate matter that provided real-time estimates of cigarette smoke intensity. Animals were monitored continuously, and analytics demonstrated particulate matter (200 μg/l of total particulate matter) and CO levels (∼1%–3%) typical of other animal exposure systems.(Basil et al., 2022; Hussain et al., 2022; Kaza et al., 2021; Raju et al., 2016) (Use approved by IACUC UAB-20232)

### Culturing of ferret and human airway epithelial cells

To obtain primary ferret tracheal epithelial cells (FTECs), freshly harvested ferret trachea were placed in DMEM media containing the antibiotics Ampho B (4 mL; 125 µg/mL), gentamicin (1.25 mL; 50 mg/mL final conc), and pen strep (10 mL). The trachea was then cut into pieces in a Biosafety Cabinet 2 under sterile conditions and incubated in a pronase (Sigma Aldrich) solution overnight at 4°C without agitation to achieve protease dissociation of the trachea. The next day, protease activity was halted using 10% FBS F12 solution. Cell pellets were harvested by centrifugation at 1000 rpm for 5 min at 4°C, resuspended with 10 mL 10% FBS F12, and filtered through a cell strainer for additional centrifugation at 1000 rpm for 5 min at 4°C. Media was then aspirated and the resulting cell pellets were dissolved in BEGM media. Cells were seeded at a density of 1x10^6^ per filter in filters coated with collagen type IV (Sigma Aldrich) and grown at the air-liquid interface (ALI) for 3-4 weeks until terminally differentiated, as described (Kaza et al., 2021).

Primary HBE cells were isolated from lung explants at the time of organ transplantation from COPD patients, healthy donors without lung disease, and healthy donors, expanded in submerged culture for one or two passages in bronchial epithelial growth medium (BEGM, Lonza, Walksville, MD), and then seeded on Transwell membranes (Corning, New York, NY) as described previously (Hussain et al., 2018; Raju et al., 2013; You et al., 2002). Cells were grown at ALI in PneumaCult-ALI media (Stem Cell Technology) for 3-4 weeks until terminally differentiated. All cells were incubated at 37^0^C and 5% CO_2_. All studies using human cells were approved by the University of Alabama at Birmingham (UAB) Institutional Review Board prior to experimentation (UAB IRB-10111015)

### Preparation of cigarette smoke extract (CSE) and treatments

CSE was generated using filtered cigarettes (1R6F: University of Kentucky, USA), absorbed on dimethyl sulfoxide (DMSO), then sterile-filtered before using in cell culture and storing at -80^0^C for further use. For all studies, well-differentiated epithelial cells were treated with 1% cigarette smoke extract or vehicle control apically (Galdi et al., 2021; Lin et al., 2020; Raju et al., 2017).

### Next-generation sequencing on Illumina platforms

The NextSeq 500 System (Illumina Inc., San Diego, CA) was used as described by the manufacturer to perform mRNA-sequencing. Briefly, the quality of the total RNA was assessed using the Agilent 2100 Bioanalyzer. RNA with an RNA Integrity Number (RIN) of 7.0 or above was used for sequencing library preparation. The Agilent SureSelect Strand Specific mRNA library kit was used as per the manufacturer’s instructions (Agilent, Santa Clara, CA). Library construction began with two rounds of polyA selection using oligo dT containing magnetic beads. The resulting mRNA was randomly fragmented with cations and heat, which was followed by first-strand synthesis using random primers with the inclusion of Actinomycin D (2.4ng/µL final concentration). Second strand cDNA production was done with standard techniques and the ends of the resulting cDNA were made blunt, A-tailed, and adaptors ligated for indexing to allow for multiplexing during sequencing. The cDNA libraries were quantitated using qPCR in a Roche LightCycler 480 with the Kapa Biosystems kit for Illumina library quantitation (Kapa Biosystems, Woburn, MA) prior to cluster generation. Cluster generation was performed according to the manufacturer’s recommendations for onboard clustering (Illumina). Paired-end 75bp sequencing runs were completed to allow for better alignment of the sequences to the reference genome.

### RNA-Seq analysis

STAR (version 2.7.0b) was used to align the raw RNA-Seq fastq reads to the ferret reference genome (Ensembl MusPutFur1.0) (Dobin et al., 2013). Following alignment, HTSeq-count (version 0.11.2) was used to count the number of reads mapping to each gene (Anders et al., 2015). Normalization and differential expression were then applied to the count files using DESeq2 (Love et al., 2014). In brief, any gene that did not have at least one read in the sum of all of the samples was removed from the analysis, leaving a total of 18,440 protein-coding genes that were relatively expressed (Hussain et al., 2022).

### Quantitative real-time PCR (qRT-PCR)

To perform qRT-PCR, RNA was first isolated from snap-frozen ferret lung tissue and cells using a Qiagen miRNeasy Mini Kit (Cat. #/ID: 217004) as per manufacturers’ protocol. Then, DNase I treatment was performed to remove DNA from the samples according to the manufacturer’s instructions (Cat. /ID:79254). Primers and probes were designed and procured from Thermo Fisher (See Supplementary list star method). qRT-PCR analysis was then carried out using a TaqMan® RNA-to-CT™ 1-Step Kit 200 reaction and Applied Biosystems Real-Time PCR system on ABI quant studio. Relative quantification of the gene of interest was calculated after normalization with the housekeeping genes, GAPDH or beta-actin, and fold change was calculated using the DDCt method (Livak and Schmittgen, 2001).

### Western Blot

Lung biopsies were isolated from healthy individuals, and smokers with and without COPD. 50mg of biopsy samples were homogenized using RIPA buffer containing 1X protease inhibitors cocktail (Thermo Fisher Scientific Cat. # 78410). The protein concentration was determined using a BCA protein assay kit (Thermo Fisher Scientific). For detection of TMPRSS2, a total of 30 µg of protein was separated on 4-12% Bis-tris SDS polyacrylamide gels (reducing) and then subjected to dry blot transfer onto nitrocellulose membranes according to the manufacturer’s instructions (Life Technologies). TMPRSS2 was detected by Western blotting using monoclonal anti-TMPRSS2 antibody (clone P5H9-A3; Millipore Sigma). Corresponding HRP anti-mouse antibody was used as secondary. Bio-Rad imagers were used for imaging and analysis (Bio-Rad). Beta-actin was used as a loading control (Cat. # T5168, Sigma Aldrich).

### H&E and AB-PAS Staining

Serial sections 5 μm thick were cut from the formalin fixed, Histogel (Fisher Scientific, Cat. # NC1095248) pre-embedded then paraffin-embedded tissue blocks and floated onto charged glass slides (Super-Frost Plus, Fisher Scientific, Pittsburgh, PA) and dried for at least 30 min at 60°C. All sections used for H&E and AB-PAS (Alcian Blue–PAS) staining were deparaffinized and hydrated using xylene (3 changes) and graded concentrations of ethanol (2 changes of 100% ETOH then 2 changes of 95% ETOH) to the tap water (H&E) or distilled water (ABPAS). For H&E staining, slides were stained in hematoxylin (STAIN HARRIS, Cardinal Health, Cat. # S7439-33) for 2 min, then differentiated in 1% acid alcohol quickly (1 dip) and bluing in ammonia water (7 dips). After hematoxylin stain, the slides were stained in eosin (stain Eosin-Y, Cardinal Health, Cat. # S7439-24) for 30 sec followed by graded concentrations of ethanol then xylene for clearing prior to the cover glass step using Permount. For AB-PAS staining the slides were pre-incubated in 3% acetic acid for 1 min then stained in Alcian Blue for 30 min. After washing in tap water then distilled water, the slides were stained in Schiff reagent (Fisher, NC9793886) for 10 min followed by reducing rinse with 0.5% sodium metabisulfite for 5 min then tap water rinse for 10 min. The slides were dehydrated through graded concentrations of ethanol then xylene for clearing prior to the cover glass step using Permount, as described (Carlson., 1997).

### Immunohistochemical staining

Serial sections 5 μm thick were cut from the formalin-fixed, paraffin-embedded tissue blocks and floated onto gelatin-coated charged glass slides (Super-Frost Plus, Fisher Scientific, Pittsburgh, PA) and dried overnight at 60°C. All sections for immunofluorescent stain were deparaffinized and hydrated using graded concentrations of ethanol to deionize water. The tissue sections were subjected to antigen retrieval by 0.01 M Tris-1mM EDTA buffer (pH 9) in a pressure cooker for 5 min (buffer preheated with the steam setting for 10 min). Following antigen retrieval, all sections were washed gently in deionized water, then transferred in to 0.05 M Tris-based solution in 0.15M NaCl with 0.1% v/v Triton-X-100, pH 7.6 (TBST). Slides were incubated with 3% Normal Horse Serum (Sigma) for 45 min at RT. All slides then were incubated at 4°C overnight with a single IF stain of ACE-2 (1:100, ab15348, Abcam) or ACE-2 (1:100, ab15348, Santa Cruz). Negative controls were produced by eliminating the primary antibodies from the diluents. After washing with TBST, slides for ACE-2 single stain and slides for ACE-2 stain were incubated with the Donkey Anti-Rabbit IgG NorthernLights™ NL557-conjugated Secondary Antibody (1:100, NL004, R&D system) and the mixture of Donkey Anti-Rabbit IgG NorthernLights™ NL557-conjugated Secondary Antibody (1:100, NL004, R&D system) and Donkey Anti-Mouse IgG NorthernLights™ NL493-conjugated Antibody (1:100, NL009, R&D system) respectively, 30 min at RT. After PBS washing, the slides were covered with cover-glasses using ProLong™ Gold Antifade Mountant with DAPI (Fisher, P36931).

### Infection studies

The SARS-CoV-2 isolate USA-WA1/2020 was obtained from BEI Resources (Cat. # NR-52281, Manassas, VA). Studies involving live viruses were conducted in the UAB BSL3 high-containment facility (Southeastern Biosafety Laboratory Alabama Birmingham; SEBLAB) with appropriate institutional biosafety approvals. For the further propagation of SARS-CoV-2, stock was passaged using Vero-E6 cells (Cat. # C1008, ATCC) and aliquoted and stored at -80^0^C until further use. Viral titers were determined by the standard Foci Forming Assay (FFU). Sequencing did not detect any mutation from the source(Campos-Gomez et al., 2023; Li et al., 2023; Pacl et al., 2021). Terminally differentiated ALI culture infection with SARS CoV-2 was conducted in the UAB SEBLAB. The apical portion of ALI culture was washed with 1X PBS to remove any mucus before infection then infected apically with 100 uL of SARS-CoV-2 viral inoculum at an MOI of 3 in all the studies prepared in EMEM media. Similarly, mock-treated cells were washed with 1X PBS followed by EMEM inoculum without any virus for 1 hr in the incubator at 370°C. At the end inoculum was removed gently using the suction pump and kept back in the incubator at 370°C with 5% CO_2_ for 72 hr.

To observe the cytopathic effect at 72 hrs post-infection, viral infection was examined by quantifying genomic and sub-genomic copy number by qPCR and immunofluorescence (IF) analysis using a SARS-CoV-2 antibody Polyclonal Anti-SARS Coronavirus antibody (Cat. # 40143-R001 from Sino Biological). SARS-CoV-2 antigen was detected in the cytoplasm of the infected cells, revealing active viral infection.

### Viral titer measurement

ALI cultures were infected with SARS-CoV-2 (MOI of 3) in Transwell plates. Mock infected cells received only the media used for preparing the SARS-CoV-2 inoculum. After 1 hr of incubation at 37^0^C with 5% CO_2_, the inoculum was removed. At each time point (days 1, 2, and 3), media from the basal chamber from mock and SARS-CoV-2-infected wells were collected and stored at negative 80^0^C. Viral production by infected ALI at each time point was measured by quantifying FFU/μL described (Gauger and Vincent, 2014). In brief, Vero-E6 cells were plated in 96-well plates at a density of 5 x 10^3^ cells/well. The next day, culture media samples collected from ALI at various time points were subjected to 10-fold serial dilutions (10^1^ to 10^6^) and inoculated onto Vero-E6 cells. The cells were incubated at 37^0^C with 5% CO_2_. After 72 hr, each inoculated well was examined for presence or absence of viral CPE and percent infected dilutions immediately above and immediately below 50% were determined. FFU was determined by ImageJ software (Schneider et al., 2012).

### Transepithelial electrical resistance (TEER)

TEER data was measured in real-time using an Epithelial colt-ohm meter (EVOM2, World Precision Inc). TEER data were collected before and after infection with SARS-CoV-2 (72 hr). Three readings were collected per well and the average was calculated.

### RNA-fluorescent in-situ hybridization (RNA-ISH)

RNA-ISH was performed using the RNAscope Multifluorescent Assay on 4% paraformaldehyde fixed 5 mm tissue section using the RNAscope reagent kit as per manufacturer’s instructions (Advance Cell Diagnostics, USA). Sections were deparaffinized with xylene for 5 min, followed by incubation with different gradients of ethanol, and then followed by incubation with hydrogen peroxide for 10 min, followed by target retrieval in the steamer for 15 min, and followed by incubation with protease plus (Advanced Cell Diagnostics) for 15 min at 40^0^C. Slides were hybridized with probes for SARS-CoV-2, ACE-2 and TMPRSS2 for 2 hr at 40°C using a HyBEZ oven and the signal was amplified by tandem incubation with Amp-1, Amp-2, Amp-3 and HRP-tagged probe (ThermoFisher Inc) for 30, 15, 30, and 15 min, respectively, at 40°C, using a HyBEZ® oven Probes were detected using Tyramide signal amplification (TSA) reaction (1:600 dilution) using an Alexa-flour labeled TSA kit (PerkinElmer) as per manufacturer’s instructions. Finally, slides were washed with wash buffer, counterstained with DAPI, and mounted with a coverslip using ProLong Gold antifade mounting media (Thermo Fisher, Rockford, IL). Images were acquired using a Nikon A1 Confocal microscope (Nikon TE2000 inverted microscope provided by the UAB Microscopy Core).

### *In situ* detection of apoptosis in cells probed with fluorescent SA-Dy546

Cells were deparaffinized in xylene and hydrated in graded ethanol series and deionized water. For immunohistochemical staining, deparaffinized and hydrated cells were washed in 0.05% v Brij-35 in PBS (pH 7.4) and immunostained as described by the manufacturer’s protocol with some modification (TACS•XL® DAB In Situ Apoptosis Detection Kit Trevigen, Cat. # 4828-30-DK). The TdT labeled tissues were detected using respective secondary antibodies conjugated with fluorescent dyes (Jackson ImmunoResearch Lab Inc., West Grove, PA). SA-Dy549 Solution (1:1000, diluted in 0.2% Triton-X 100 /PBS) was incubated for 30 min at room temperature. Where indicated, the sections were stained with 4’,6-diamidino-2-phenylindole (DAPI) containing Fluormount-G (SouthernBiotech, Birmingham, AL) to visualize nuclei. Immunofluorescent images were captured with the BZX700 Microscopy system (Keyence Corp., Japan) (Hussain et al., 2018).

### ACE-2 antisense oligonucleotide (ASO) and camostat mesylate treatment

ASOs targeting ACE-2 were designed and synthesized through Integrated DNA technology (IDT)) at a concentration of 1000 µM, prepared in sterile PBS, and evaluated in primary FTECs grown at ALI. To establish baseline efficacy and optimal dose, cells (1 x 10^6^ cells/mL) were treated for 1 week with 5 different ACE-2 ASOs based on different regions of nucleotide sequence at concentrations of 10 µM, 20 µM, 100 µM, after which knockdown was assessed using qRT-PCR (Supplemental Figure 2A). Control ASO and untreated conditions were included. For studies evaluating efficacy in the context of smoking and SARS-CoV-2, cells were treated for 7 days with the most efficacious ACE-2 ASO at an optimized concentration of 20 µM (Supplemental Figure 2B) or control ASO. Cells were then exposed to vehicle or whole CSE produced in DMSO as solvent due to its characterized properties in our lab (Galdi et al., 2021) for 48 hr, and finally infected with SARS-CoV-2 at an MOI of 3 for a 72-hr infection period. For studies with combination ACE-2 and camostat mesylate treatment, FTECs (5 x 10^5^ cells/mL) were treated with ACE-2 ASO or control ASO for 7 days, with acute addition of CSE or vehicle on day 7 and 8. On day 9, cells were washed and then treated with camostat mesylate (100 µM) for 2 hr prior to inoculation with SARS-CoV-2. Cells were harvested on day 12 for analysis.

### Statistical analyses

Quantitative metrics were analyzed using GraphPad Prism (GraphPad Software, La Jolla, CA, USA), deploying unpaired t-tests, one-way ANOVA, or two-way ANOVA with proper post-hoc tests as appropriate. Unless noted otherwise, data are presented as Mean ± SEM. P-values < 0.05 were considered statistically significant.

## Acknowledgments

These studies were supported by NIH National Heart, Lung, and Blood Institute (NHLBI) Grant R35HL135816 (to S.M. R) and National Institute of Diabetes and Digestive and Kidney Diseases Grant P30 DK072482 (to S.M. R). Cystic fibrosis foundation, Funded in part by a grant from the Alpha-1 Foundation 919960 (To S.S.H).

## Author Contributions

S.S.H and S.M.R conceived of and designed the research; S.S.H, performed the experiments, analyzed data, and interpreted results; S.S.H, E.F.L, J.L.T, S.E.P, Q.L, J.C.G, J.P.L, M.M, K.S.H, and S.M.R formal analysis; S.S.H, E.F.L, and S.M.R prepared figures; S.S.H, E.F.L, and S.M.R wrote the manuscript; S.M.R supervised the project; all authors had an opportunity to edit the manuscript and approved of its submission. We acknowledge the Southern Biosafety Laboratory at the University of Alabama at Birmingham (SEBLAB) and the NIAID-supported laboratory (UC6 AI058599) for their work related to BSL-3 containment.

---

**S. Figure 1.**
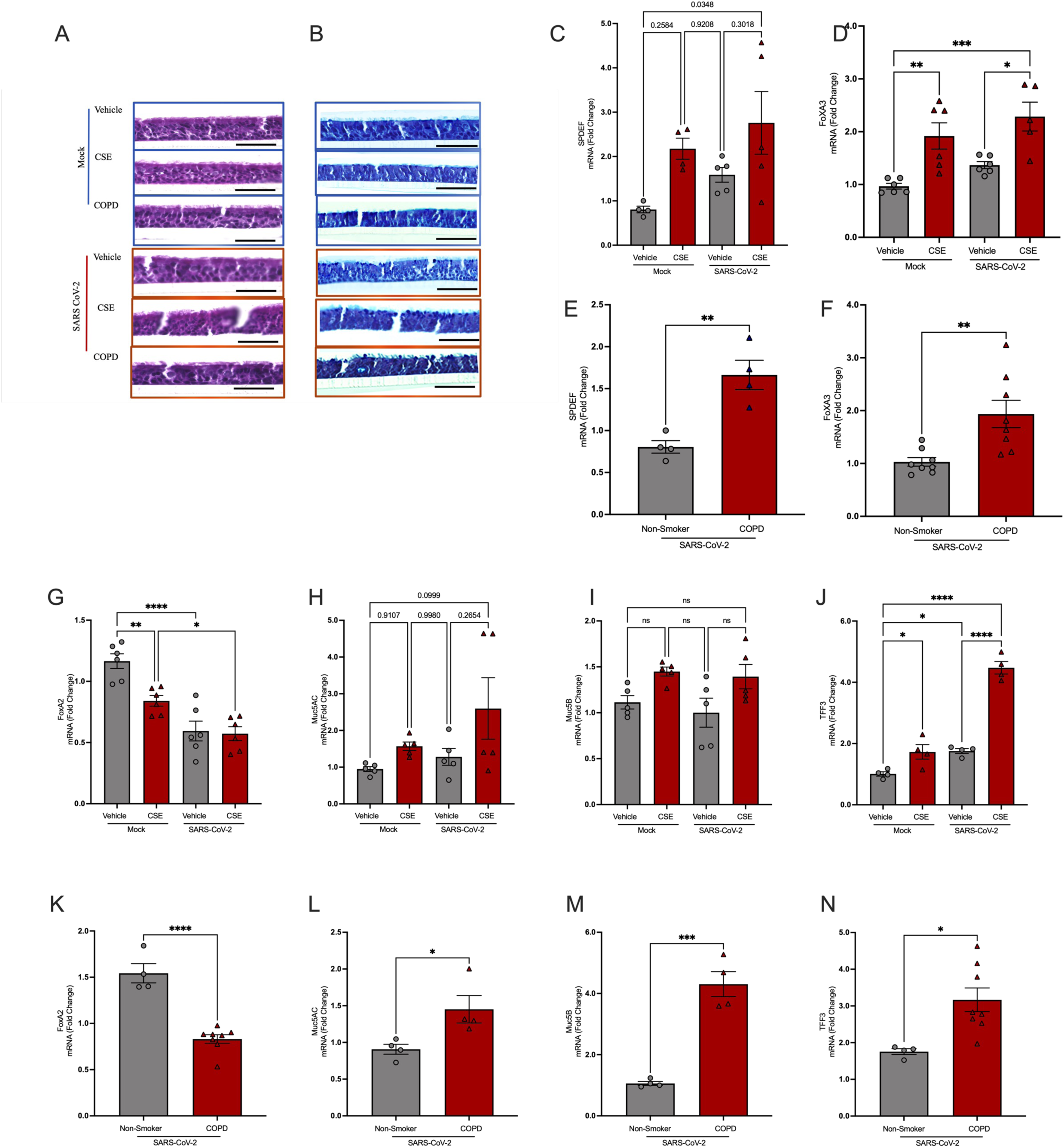
Cigarette smoking promotes SARS-CoV-2-induced goblet cell hyperplasia and mucin production in CSE-exposed normal HBE cells and in COPD HBE cells. Representative images of (A) hematoxylin and eosin (H&E) and (B) Alcian blue/periodic acid-Schiff (AB/PAS) staining of vehicle-treated, cigarette smoke extract (CSE)-treated, and COPD human bronchial epithelial (HBE) cells infected with SARS-CoV-2 or mock infection. Scale bar, 50 μm. (C-M) qRT-PCR quantification of transcription factor and secretory protein mRNA expression in each group. *P<0.05, **P<0.01, ***P<0.001, ****P<0.0001.

**S. Figure 2.**
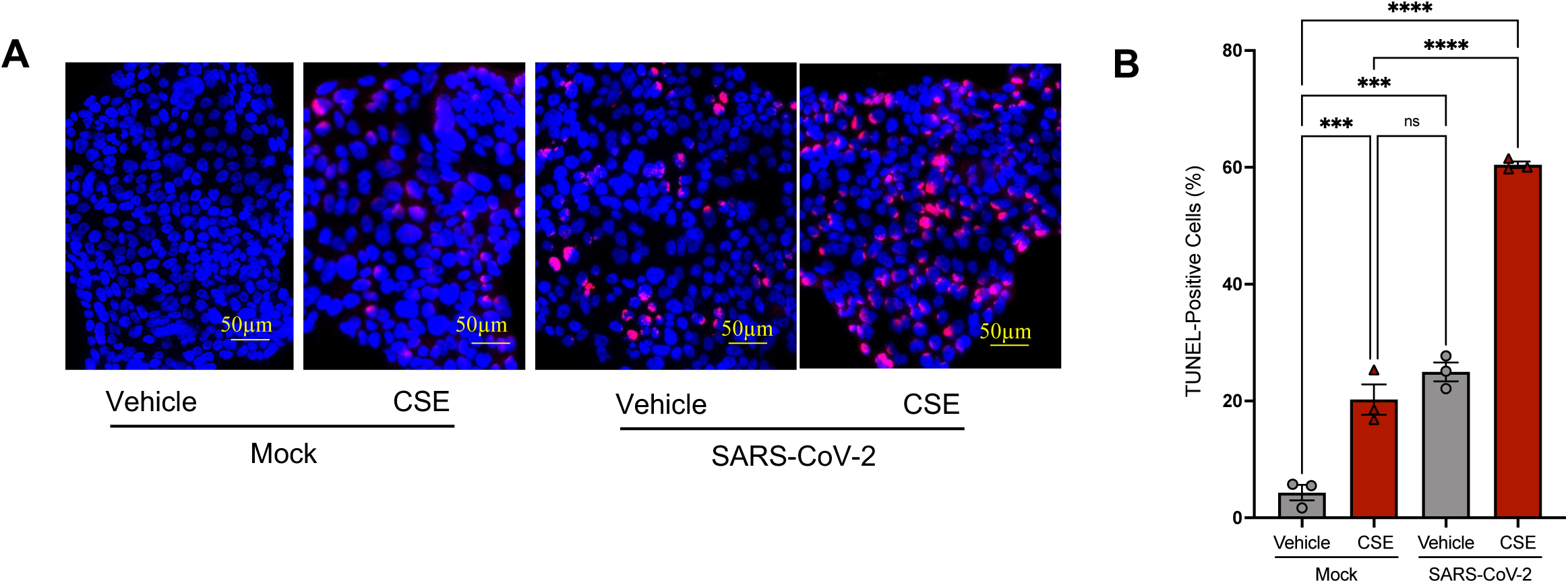
Apoptosis. (A) Representative images and (B) quantification of TUNEL staining in human bronchial epithelial (HBE) cells treated with cigarette smoke extract (CSE) or vehicle control for 48 hr followed by 72 hr infection with SARS-CoV-2 or mock infection. TUNEL positivity (pink) and DAPI-stained nuclei (blue). N=3-5 per well per group. Scale bar, 50μM. ***P<0.001, ****P<0.0001

**S. Figure 3.**
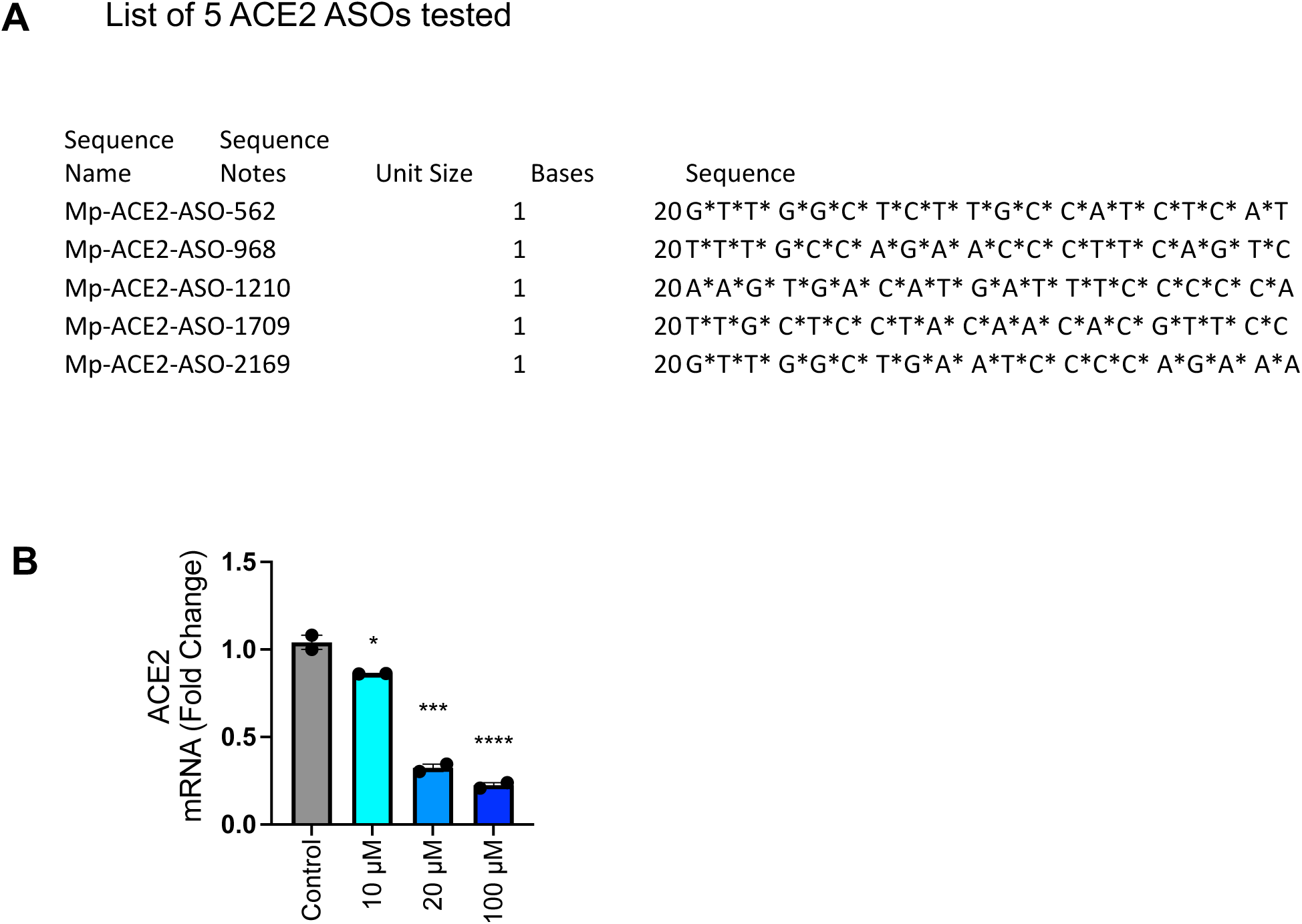
A) List of ACE-2 specific ASOs and sequence used for preliminary testing of ACE-2 knockdown in FTEC (B) Dose-response evaluation of ACE2 mRNA knockdown in ferret tracheal epithelial cells (FTECs) treated with control ASO or different concentrations of ACE2 ASO for an alternate day for five days.

**S. Figure 4.**
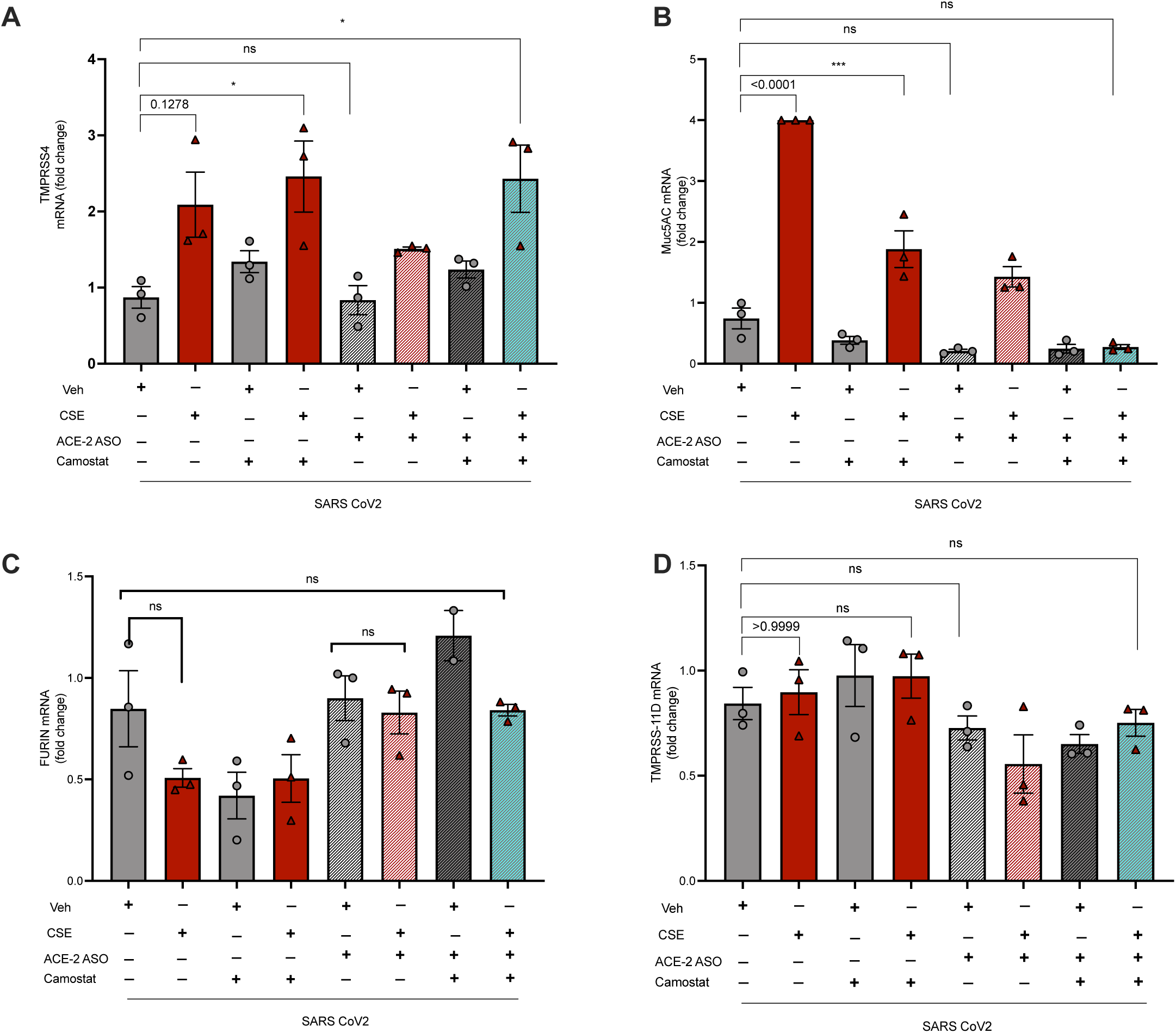
(B) qRT-PCR quantification of MUC5AC shows attenuation of the MUCIN gene following ACE-2 knockdown. Other genes associated with SARS-CoV-2, such as TMPRSS4 (A), FURIN (C), and TMPRSS1D (D), did not exhibit significant changes in mRNA expression across the groups. *P<0.05, **P<0.01, ***P<0.001, ****P<0.0001

